# A susceptible mouse model for Zika virus infection

**DOI:** 10.1101/042358

**Authors:** S.D. Dowall, V.A. Graham, E. Rayner, B Atkinson, G. Hall, R. Watson, A. Bosworth, L. Bonney, S Kitchen, R. Hewson

## Abstract

Zika virus (ZIKV) is a mosquito-borne pathogen which has recently spread beyond Africa and into Pacific and South American regions. Despite first being detected in 1947, very little information is known about the virus and its spread has been associated with increases in Guillain-Barre syndrome and microcephaly. There are currently no known vaccines or antivirals against ZIKV infection. Progress in assessing interventions will require the development of animal models to test efficacies; however, there are only limited reports on *in vivo* studies. The only susceptible murine models have involved intracerebral inoculations or juvenile animals, which do not replicate natural infection. Our report has studied the effect of ZIKV infection in type-I interferon receptor deficient (A129) mice and the parent strain (129Sv/Ev) after subcutaneous challenge in the lower leg to mimic a mosquito bite. A129 mice developed severe symptoms with viral RNA being detected widespread in the blood, brain, spleen, liver and ovaries. Histological changes were also striking in these animals. 129Sv/Ev mice developed no clinical symptoms or histological changes, despite viral RNA being detectable in the blood, spleen and ovaries, albeit at lower levels to those seen in A129 mice. Our results identify A129 mice as being highly susceptible to ZIKV and thus a suitable small animal model for the testing of vaccines and antivirals which are urgently required.

**Author summary:** Since first being recognised in 1947, Zika virus (ZIKV) has mainly been a mild illness with symptoms including a limited fever and rash. In 2007 the virus spread from Africa into French Polynesia and then onwards across Pacific regions and into South America. In these new regions, ZIKV has been associated with more severe clinical conditions including Gullain-Barre syndrome and microcephaly. There are no currently approved antivirals or vaccines available with proven activity against ZIKV, and the World Health Organisation declared the spread of ZIKV as a Public Health Emergency of International Concern. Here, we have used a mouse strain with a deficiency in the type-I interferon receptor (A129) and shown that they are susceptible to ZIKV infection after inoculation that closely resembles the natural route via mosquito bite. Although A129 mice are deficient in the innate interferon response, they retain their adaptive immunity and thus have successfully been used as suitable models for the testing of vaccinations and antivirals. Our study provides details on a suitable model for the testing of future interventions against ZIKV.

## Introduction

Zika virus (ZIKV) is a mosquito-borne flavivirus first isolated from a sentinel rhesus macaque placed in the Zika forest in Uganda in 1947; thereafter, ZIKV was isolated from a pool of *Aedes africanus* mosquitoes from the same forested area [1]. During the 1960s-1980s evidence for ZIKV infection was detected in several African and Asian countries including Nigeria [2], Pakistan [3], Malaysia [4] and Indonesia [5]. Until an outbreak in 2007 on Yap Island, Micronesia, no transmission of Zika virus had been reported outside of Africa and Asia and only 14 cases of human ZIKV disease had been previously documented [6]. No further transmission was identified until 2013 when French Polynesia reported autochthonous cases [7] which led to an epidemic in which an estimated 28,000 people sought medical advice (11% of the population) [8]. The virus then spread rapidly throughout the Pacific region [9] and there was an imported case into the UK, a male tourist who visited the Cook Islands in 2014. Interestingly, this case showed evidence of ZIKV RNA in semen two months post onset of symptoms and underlined the prolonged potential for sexual transmission of ZIKV [10]. In 2015, ZIKV was reported in Brazil [11]. Whilst it is not known how it became established in this previously non-endemic region, one theory presented by Brazilian researchers is that it entered the country during the 2014 football World Cup [12], another is that it was introduced during the Va’a World Sprint Championship canoe race held in Rio de Janeiro, in 2014 [13]. The latter is supported by phylogenetic studies [14] and also that no ZIKV-endemic Pacific countries competed in the football World Cup.

The expansion of ZIKV endemicity follows closely that of chikungunya virus, which now circulates in most inhabited continents where competent mosquito vectors are extant, and has become a global public health problem in the past decade [15]. At the time of writing, the number of countries reporting autochthonous transmission of ZIKV is increasing with the possibility of further spread into non-endemic areas including North America[16]. The first importation of a ZIKV infected traveller into the US during the ongoing outbreak was reported in January 2016 [17]. On 1^st^ February, 2016, the World Health Organisation declared ZIKV to be a “Public Health Emergency of International Concern” [18].

Recently, ZIKV infection has been associated with Guillain-Barre syndrome (GBS), an autoimmune disease that causes acute or subacute flaccid paralysis [19, 20]. In December 2013, a patient from French Polynesia presented with GBS a week after confirmation of acute ZIKV infection [21]. Subsequent GBS cases were confirmed and correlated temporally with the ZIKV outbreak [9].The incidence rates of GBS during the ZIKV outbreak were approximately 20-fold higher than expected given both the size of the French Polynesian population and the established incidence rates of GBS (1-2 per 100,000 population per year) [22]. Whilst the recent temporal and spatial association between the ZIKV outbreak and the increase in cases of GBS implies a potential link, at present there are no data to confirm ZIKV as the antigenic stimulus predisposing individuals to this autoimmune disease [9].

Recent widespread attention has been given to increases in the cases of microcephaly and its coincidence with the ZIKV epidemic in South America. Over the past five years, the incidence of microcephaly was between 130 and 170 cases annually, but in the first nine months of 2015 this figure roughly doubled [23]. In the last three months of 2015, over 2400 further cases were reported [23]. This coincides with the rapid spread of ZIKV which was first confirmed in Brazil in May 2015 [11], indeed, ZIKV has been shown to be present in abnormal foetal brain tissue linked to microcephaly [48, 49]. However, clear scientific evidence of this aetiology has yet to emerge. Several other infections and toxins are known to cause foetal brain insults and teratogenic effects [24], including: West Nile virus [25, 26]; rubella virus [27]; cytomegalovirus [28]; herpes simplex virus type 1 [29]; varicella zoster virus [30]; human immunodeficiency virus [31]; lymphocytic choriomeningitis virus [32]; and mosquito larvicide [33]. Furthermore the diagnosis of ZIKV infection is not straightforward [34]; the opportunity to detect viral RNA in serum is often missed and serological techniques show extensive cross reactivity between antibodies triggered by different flavivirus infections or past vaccination. Thus there is an immediate and urgent need for research work into ZIKV with a priority on experimental studies *in vivo*.

A small animal disease model would advance knowledge about this pathogen, and open avenues for further research including studies of ZIKV vertical transmission, whether infection has direct or indirect effects on neural development [35] as well as providing a means to assess novel interventions, determining pathogenicity of different strains, and the effects of mutations. Mouse models reported previously have relied on the use of juvenile animals and/or intracerebral inoculations [1, 2, 36-44]. In this work we have developed an adult model and simulated a natural route of infection. We believe this model will enable a wider range of studies to advance our knowledge of ZIKV.

## Methods

### Virus stocks

ZIKV (strain MP1751, isolated in 1962 from pools of *Aedes Africanus* [40]) was obtained from the National Collection of Pathogenic Viruses, UK. Virus was cultivated in Vero cells obtained from the European Collection of Cell Cultures, UK, with Dulbecco’s Modified Eagle Medium containing GlutaMAX (Invitrogen, UK) and supplemented with 2% heat-inactivated foetal bovine serum (Sigma, UK). Infectious virus titres were determined by plaque assay on Vero cells incubated for six days and using a 1% carboxymethyl cellulose overlay.

### Mice

Mice deficient in the IFN-α/β receptor (A129) and congenic control mice (129Sv/Ev) were obtained from an established breeding colony approved by the UK Home Office (B&K Universal, UK). Female mice aged 5-6 weeks were used for all studies. All procedures were undertaken according to the United Kingdom Animals (Scientific Procedures) Act 1986. These studies were approved by the ethical review process of Public Health England, Porton Down, UK and the Home Office, UK via an Establishment Licence (PEL PCD 70/1707) and project licence (30/3147).

### Pathogenicity studies

Virus inocula containing 10^6^ plaque-forming units (pfu) of ZIKV diluted in phosphate buffered saline (PBS) were administered subcutaneously in volumes of 40μL into each of the right and left hind legs just above the ankle to 12 A129 and 12 129Sv/Ev mice. Control groups of a further three animals of each mouse strain were inoculated with 40 μL PBS in each of the hind legs. Mice were monitored three times each day for clinical signs of disease and a numerical score assigned at each observation, (as per: 0 normal; 2 ruffled fur; 3 lethargy, pinched, hunched, wasp waisted; 5 laboured breathing, rapid breathing, inactive, neurological; and 10 immobile). Temperatures were recorded by an indwelling temperature chip and weights were also recorded daily. A set of humane clinical end points were defined by veterinary staff as a 20% weight loss, or 10% weight loss and a clinical symptom, which mandated euthanasia. At 3 and 7 days post-challenge, a group of four animals from each of the A129 and 129Sv/Ev ZIKV-challenged groups were scheduled to be culled to assess local responses. At necropsy, samples of spleen, liver, brain and ovary were collected and immediately frozen at −80°C for virological analysis or inserted into pots containing 10% neutral buffered saline for microscopic analysis. Blood was collected into RNAprotect tubes (Qiagen, UK) for viral load testing.

### Real-time RT-PCR assay

Tissue samples were weighed and homogenised into PBS using ceramic beads and an automated homogeniser (Precellys, UK) using six 5 second cycles of 6500 rpm with a 30 second gap. 200 μL of tissue homogenate or blood solution was transferred to 600 μL RLT buffer (Qiagen, UK) for RNA extraction using the RNeasy Mini extraction kit (Qiagen, UK) with an initial step of passing samples through a QIAshredder (Qiagen, UK).

A ZIKV specific real-time RT-PCR assay was utilised for the detection of viral RNA from subject animals. The primer and probe sequences were adopted from a published method [45] with in-house optimisation and validation performed to provide optimal mastermix and cycling conditions (data not shown). Real-time RT-PCR was performed using the SuperScript III Platinum One-step qRT-PCR kit (Life Technologies, UK). The final mastermix (15μL) comprised 10μL of 2x Reaction Mix, 1.2 μL of PCR-grade water, 0.2μL of 50mM MgSO_4_, 1μl of each primer (ZIKV 1086 and ZIKV 1162c both at 18μM working concentration), 0.8μL of probe (ZIKV 1107-FAM at 25μM working concentration) and 0.8μL of SSIII enzyme mix. 5μL of template RNA was added to the mastermix in order to give a final reaction volume of 20μL The cycling conditions used were 50°C for 10 minutes, 95°C for 2 minutes, followed by 45 cycles of 95°C for 10 seconds and 60°C for 40 seconds (with quantification analysis of fluorescence performed at the end of each 60°C step), and a final cooling step of 40°C for 30 seconds. Reactions were run and analysed on the 7500 Fast platform (Life Technologies, UK) using 7500 software version 2.0.6.

Quantification of viral load in samples was performed using a dilution series of quantified RNA oligonucleotide (Integrated DNA Technologies). The oligonucleotide comprised the 77 bases of ZIKV RNA targeted by the assay based on GenBank accession AY632535.2 and was synthesised to a scale of 250 nmole with HPLC purification. This RNA oligonucleotide was quantified using Nanodrop technology (Thermo Scientific, UK) with a 10-fold dilution series of quantified material used to assess viral load in subject samples. Eight dilutions of quantified ZIKV RNA oligonucleotide control material were used to assess the limit of detection ranging from 1,000,000 copies per reaction to 0.1 copies per reaction; all dilutions were run in triplicate. The limit of detection for this assay was assessed to be 10 genome copies per PCR reaction with 100% detection for this and all higher concentrations. All samples for 1 copy per reaction and 0.1 copies per reaction were negative. In-run analysis indicated the reaction efficiency to be 101% with a slope of −3.29, a Y intercept of 40.17 cycles and an R2 value of 0.998.

### Histology

Tissue samples were fixed in 10% neutral buffered formalin for 48 hours and processed routinely to paraffin wax. Sections were cut at 3-5 μm, stained with haematoxylin and eosin (H&E) and examined microscopically. Lesions referable to infection were scored subjectively using the following scale: within normal limits (WNL), minimal, mild, moderate and marked. The pathologist was blinded to the groups in order to prevent bias.

## Results

### A129 mice are susceptible to challenge with ZIKV, whilst 129Sv/Ev mice are resistant and show no clinical evidence of infection

Groups of 5-6 week old mice either lacking receptors for IFN-α/β (A129) or from the wild-type strain (129Sv/Ev) were subcutaneously inoculated with 10^6^ pfu ZIKV in the lower leg. Survival analysis was compared between the two ZIKV-infected mouse strains and mock-infected animals which received PBS only, and demonstrated that all A129 animals met humane clinical endpoints 6 days after challenge with ZIKV (Fig. 1A). Wild-type 129Sv/Ev mice all survived the 14 day length of the study, as did the control animals. When weights were compared between groups, the ZIKV-challenged A129 mice started to lose weight rapidly after day 3 post-challenge, whereas the other groups all demonstrated a gradual increase over the course of the study indicating that animals were healthy (Fig. 1B). Similarly, the temperatures of the A129 mice showed differences compared to the other groups, with a gradual increase until day 4 post-challenge and then a rapid decrease (Fig. 1C). The A129 ZIKV-challenged mice were the only ones which exhibited signs of disease post-challenge, with signs first being noted on day 5 and increasing until humane endpoints were met on day 6 post-challenge (Fig. 1D). None of the other groups had any clinical signs recorded for the duration of the study.

**Figure 1.**
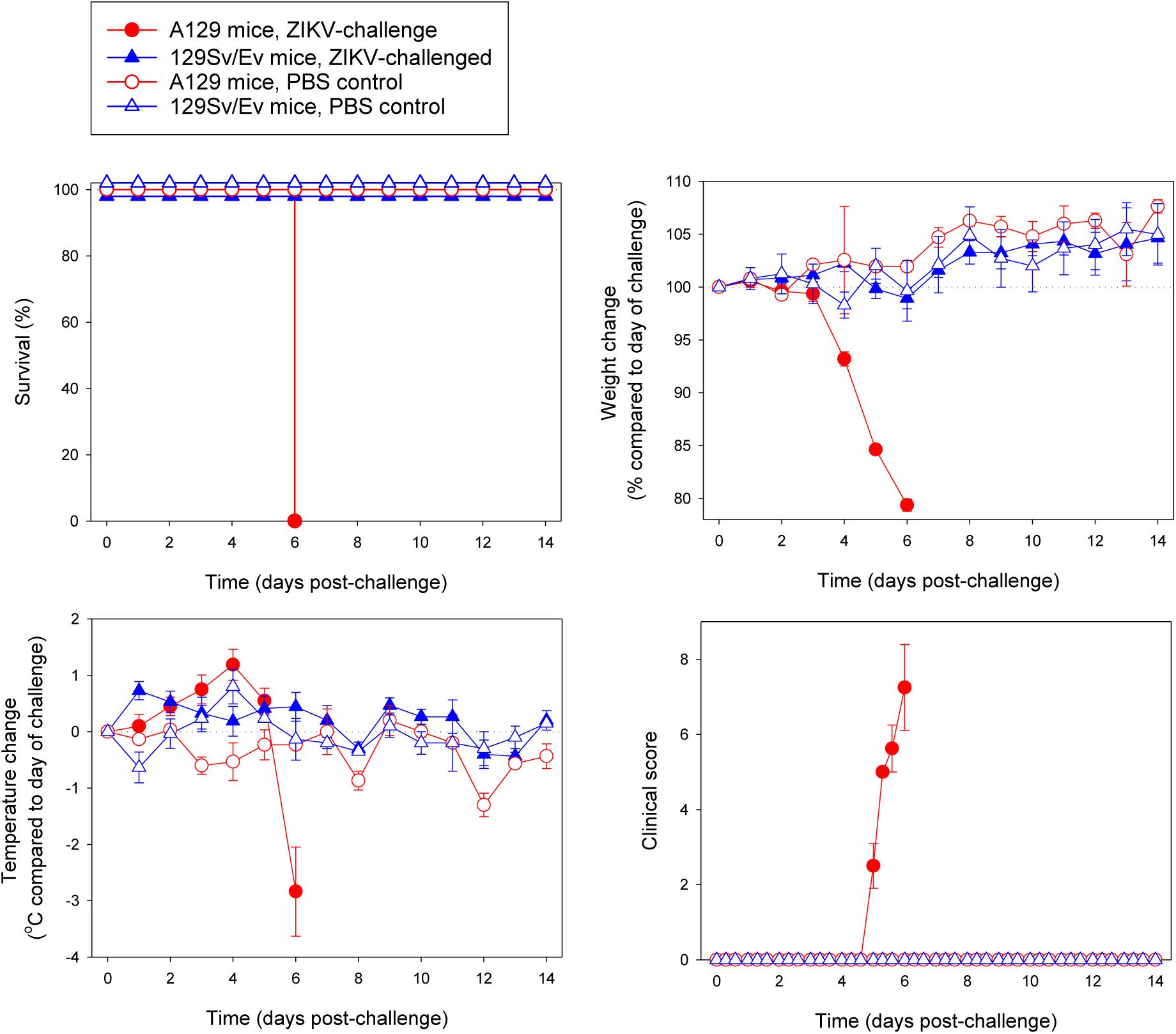
ZIKV is virulent to IFN-α/β receptor deficient mice (A129) whilst wild-type mice (129Sv/Ev) do not exhibit signs of disease. A129 (closed circle) and 129Sv/Ev (closed triangle) mice were subcutaneously inoculated in the lower leg with 10^6^ pfu ZIKV and compared to PBS mock-infected A129 (open circle) and 129Sv/Ev (open triangle) groups. **(A)** Survival analysis of groups presented as Kaplan-Meier survival curves. **(B)** Weight changes compared to the day of challenge. **(C)** Temperature changes compared to the day of challenge. **(D)** Clinical scores of animals. Graphs B-D show the mean values with errors bars denoting standard error. Group sizes were: n=8 (A129, ZIKV-challenged); n=8 to day 7 and n=4 from days 8-14 (129Sv/Ev, ZIKV-challenged); and n=3 (A129 and 129Sv/Ev, PBS mock-challenged).

### ZIKV-specific viral RNA was widespread in A129 challenged mice but at lower levels and only in certain tissues in wild-type 129Sv/Ev mice

Viral RNA levels were quantified in the blood and tissues (brain, ovary, spleen and liver) of ZIKV-challenged A129 and 129Sv/Ev mice culled at day 3 post-challenge and days 6 (A129) or 7 (129Sv/Ev) post-challenge (Fig. 2). Results demonstrated that in A129 mice, the viral RNA was detected in all of the samples at day 3 post-challenge, with levels highest in the spleen. By day 6 post-challenge, the levels remained high but were reduced in all of the samples compared to day 3 apart from the brain. For 129Sv/Ev mice, viral RNA was only detected in the blood, ovary and spleen at day 3 post-challenge, and at day 7 post-challenge was still within the organs but was undetected in the circulation. The levels of viral RNA in 129Sv/Ev mice were orders of magnitude lower than those found in A129 mice. Samples from PBS mock-challenged animals were consistently negative for viral RNA (data not shown).

**Figure 2.**
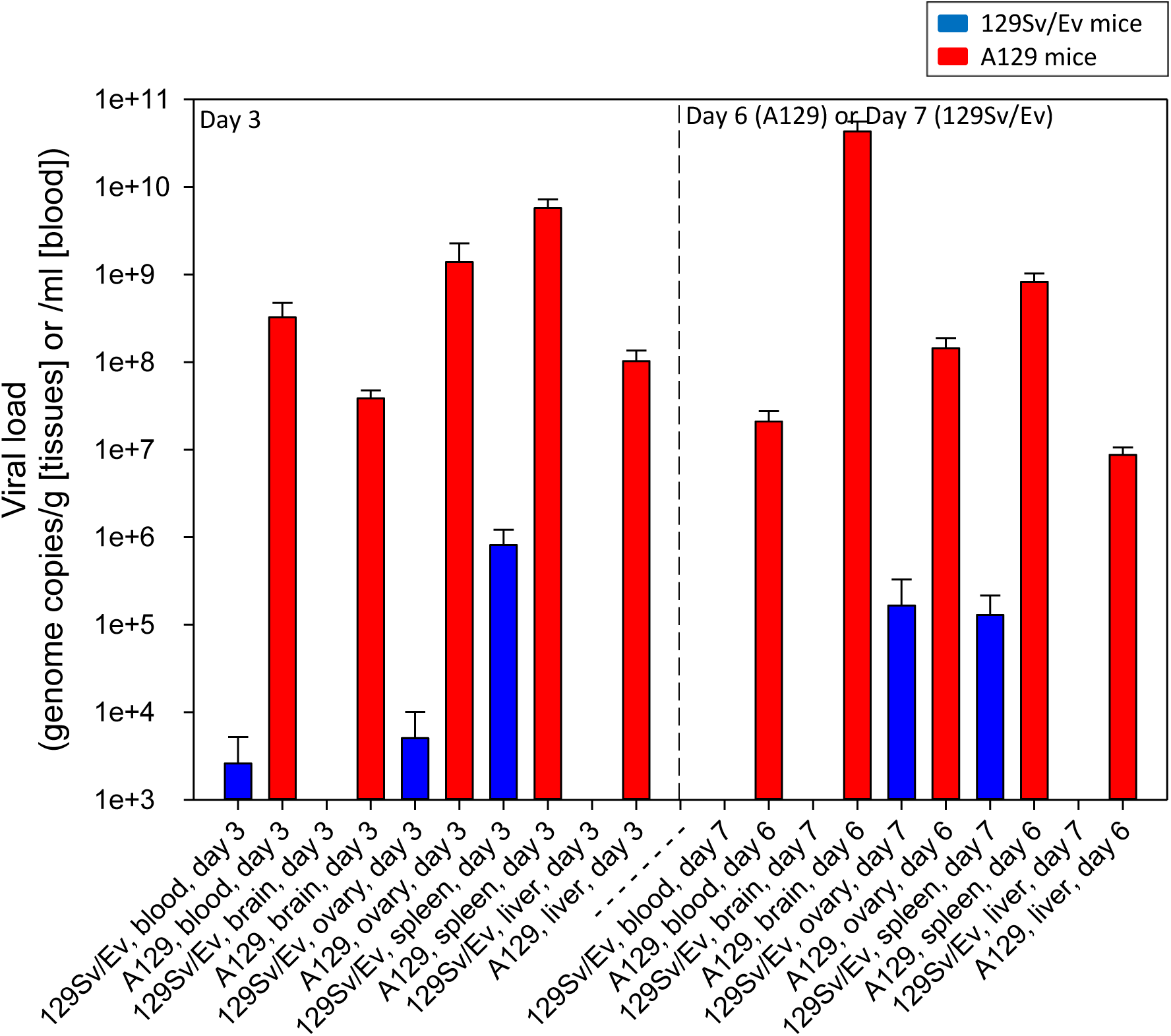
Levels of ZIKV viral RNA in blood, brain, ovary, spleen and liver samples in IFN-α/β receptor-deficient (A129) and wild-type (129Sv/Ev) mice post-challenge. Mice were subcutaneously challenged in the lower leg with 10^6^ pfu ZIKV. At day 3 and day 6 (A129) or 7 (129Sv/Ev) mice were culled to assess viral load within the circulation and at local sites. Results are denoted as the genome equivalent per ml of blood or per g of tissue. Bars show mean viral RNA levels with error bars denoting standard error. Group sizes were: n=4 (A129 and 129Sv/Ev, day 3 post-challenge); n=8 (A129, day 6 post-challenge); and n=4 (129Sv/Ev, day 7 post-challenge).

## Pathology

Inflammatory and degenerative changes were observed in the brain of A129 mice challenged with ZIKV. These comprised widespread nuclear fragments, scattered diffusely throughout the grey and white matter (Fig 3A). Perivascular cuffing of vessels was observed in the parenchyma and meninges by mononuclear cells, many having the morphology of lymphocytes (Fig 3B). Varying numbers of polymorphonuclear cells (PMNs) were noted in the grey and white matter, often near blood vessels (Fig 3C). Partially degenerated cells having hyper-eosinophilic cytoplasm and irregularly, shaped, partially condensed nuclei were noted amongst the neurons of the hippocampus (Fig. 3D).

**Figure 3.**
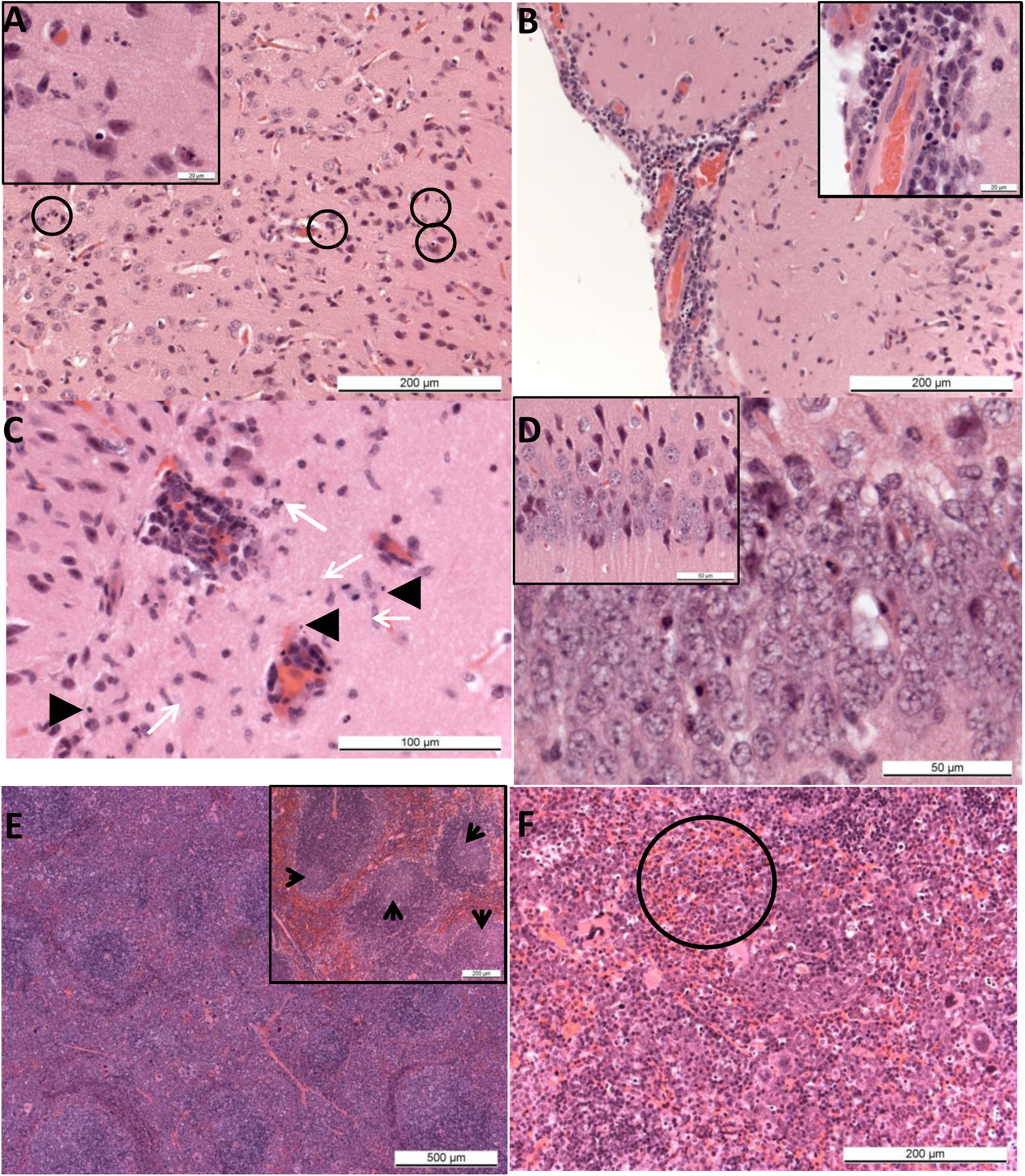
Pathological findings in ZIKV challenged A129 mice. (A) Brain. Animal 23842 (A129, ZIKV-challenged, day 6). Scattered, nuclear fragment in the neuropil of the cerebral cortex (circled). Inset, higher magnification of nuclear debris. (B) Brain. Animal 23811 (A129, ZIKV-challenged, day 6). Lymphocyte infiltration of meninges and perivascular cuffing of meningeal vessels surrounding the cerebral cortex. Inset, degenerate and fragmented nuclei. (C) Brain. Animal 23962 (A129, ZIKV-challenged, day 6). Prominent, perivascular cuffing of small vessels in the neuropil, with infiltration of polymorphonuclear leukocytes (arrows) and presence of scattered, nuclear fragments (arrowheads). H&E. (D) Brain. Animal 23842 (A129, ZIKV-challenged, day 6). Scattered, shrunken, hyper-eosinophilic cells (arrows) amongst the neurons of the hippocampus. Inset, Animal 23798 (129Sv/Ev, PBS, day 14). Normal neurons. (E) Spleen. Animal 23800 (A129, ZIKV-challenged, day 6). Poorly defined white pulp with large, irregular germinal centres. Inset, Animal 23798 (129Sv/Ev, PBS, day 14). Well defined germinal centres within the white pulp (F) Spleen, red pulp. Animal 23800 (A129, ZIKV-challenged, day 6). Numerous, mature neutrophils within the red pulp (circled). (F) Liver. Animal 23836 (A129, ZIKV-challenged, day 6). Small foci of extra-medullary haematopoiesis scattered randomly throughout the parenchyma.

Numerous small well-defined germinal centres with apoptotic bodies and mitotic figures were noted in the splenic white pulp of control animals. In the spleen of challenged A129 mice large, poorly defined germinal centres were observed in the white pulp, together with numerous apoptotic bodies and a prominent depletion in the number of mature lymphocytes, (Fig 3E). Prominent extra-medullary haematopoiesis as well as numerous mature PMNs were present in the red pulp sinuses (Fig 3F). Small foci of extra-medullary haematopoiesis were also observed in the liver. The ovaries were normal.

Lesions referable to Zika virus infection were not observed in the brain, spleen, liver or ovaries of the wild-type 129Sv/Ev mice or the control groups (Table 1).

**Table 1.**
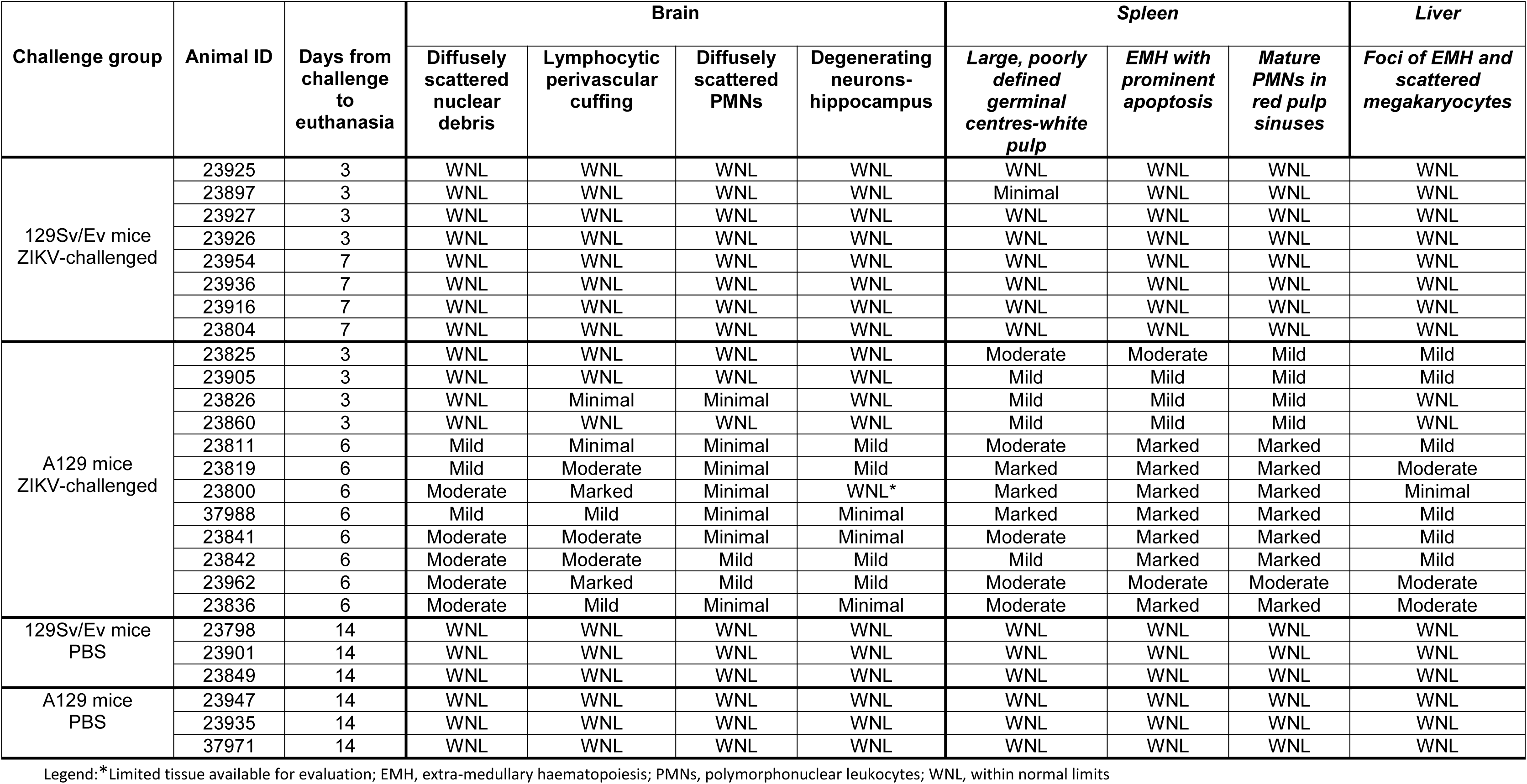
Histology changes observed in IFN-α/β receptor deficient (A129) and wild-type mice (129Sv/Ev) challenged with ZIKV or mock-challenged with PBS

| Challenge group | Animal ID | Days from challenge to euthanasia | Brain |  |  |  | Spleen |  |  | Liver |
| --- | --- | --- | --- | --- | --- | --- | --- | --- | --- | --- |
|  |  |  | Diffusely scattered nuclear debris | Lymphocytic perivascular cuffing | Diffusely scattered PMNs | Degenerating neurons-hippocampus | Large, poorly defined germinal centres-white pulp | EMH with prominent apoptosis | Mature PMNs in red pulp sinuses | Foci of EMH and scattered megakaryocytes |
| 129Sv/Ev mice<br>ZIKV-challenged | 23925 | 3 | WNL | WNL | WNL | WNL | WNL | WNL | WNL | WNL |
|  | 23897 | 3 | WNL | WNL | WNL | WNL | Minimal | WNL | WNL | WNL |
|  | 23927 | 3 | WNL | WNL | WNL | WNL | WNL | WNL | WNL | WNL |
|  | 23926 | 3 | WNL | WNL | WNL | WNL | WNL | WNL | WNL | WNL |
|  | 23954 | 7 | WNL | WNL | WNL | WNL | WNL | WNL | WNL | WNL |
|  | 23936 | 7 | WNL | WNL | WNL | WNL | WNL | WNL | WNL | WNL |
|  | 23916 | 7 | WNL | WNL | WNL | WNL | WNL | WNL | WNL | WNL |
|  | 23804 | 7 | WNL | WNL | WNL | WNL | WNL | WNL | WNL | WNL |
| A129 mice<br>ZIKV-challenged | 23825 | 3 | WNL | WNL | WNL | WNL | Moderate | Moderate | Mild | Mild |
|  | 23905 | 3 | WNL | WNL | WNL | WNL | Mild | Mild | Mild | Mild |
|  | 23826 | 3 | WNL | Minimal | Minimal | WNL | Mild | Mild | Mild | WNL |
|  | 23860 | 3 | WNL | WNL | WNL | WNL | Mild | Mild | Mild | WNL |
|  | 23811 | 6 | Mild | Minimal | Minimal | Mild | Moderate | Marked | Marked | Mild |
|  | 23819 | 6 | Mild | Moderate | Minimal | Mild | Marked | Marked | Marked | Moderate |
|  | 23800 | 6 | Moderate | Marked | Minimal | WNL* | Marked | Marked | Marked | Minimal |
|  | 37988 | 6 | Mild | Mild | Minimal | Minimal | Marked | Marked | Marked | Mild |
|  | 23841 | 6 | Moderate | Moderate | Minimal | Minimal | Moderate | Marked | Marked | Mild |
|  | 23842 | 6 | Moderate | Moderate | Mild | Mild | Mild | Marked | Marked | Mild |
|  | 23962 | 6 | Moderate | Marked | Mild | Mild | Moderate | Moderate | Moderate | Moderate |
|  | 23836 | 6 | Moderate | Mild | Minimal | Minimal | Moderate | Marked | Marked | Moderate |
| 129Sv/Ev mice<br>PBS | 23798 | 14 | WNL | WNL | WNL | WNL | WNL | WNL | WNL | WNL |
|  | 23901 | 14 | WNL | WNL | WNL | WNL | WNL | WNL | WNL | WNL |
|  | 23849 | 14 | WNL | WNL | WNL | WNL | WNL | WNL | WNL | WNL |
| A129 mice<br>PBS | 23947 | 14 | WNL | WNL | WNL | WNL | WNL | WNL | WNL | WNL |
|  | 23935 | 14 | WNL | WNL | WNL | WNL | WNL | WNL | WNL | WNL |
|  | 37971 | 14 | WNL | WNL | WNL | WNL | WNL | WNL | WNL | WNL |

Legend: \*Limited tissue available for evaluation; EMH, extra-medullary haematopoiesis; PMNs, polymorphonuclear leukocytes; WNL, within normal limits

## Discussion

At the time of writing, our studies provide the first evidence for a susceptible adult mouse model for ZIKV infection after shallow inoculation of virus below the skin surface at one of the body’s extremities, mimicking natural infection via mosquito bite [46]. Additionally, the virus used in our studies did not need adapting to cause lethality, unlike earlier studies where *in vivo* passaging of virus was needed to strengthen effects [37]. The concentration of 10^6^ pfu used for inoculation used was within the limits of studies with another mosquito-borne flavivirus, West Nile virus, which demonstrated a viral dose 10^4^-10^6^ pfu per mosquito bite [47].

Throughout the course of infection, animals were monitored regularly for clinical signs of disease, including changes in weight and temperature fluctuations. Only the A129 mice with a knockout in their type-I interferon receptor showed evidence of disease. Although fever and general malaise are typical of human ZIKV infection [48], similar observations were not seen in infected mice. The severe weight loss, hypothermia and clinical signs that were recorded however, meant that ZIKV-challenged A129 mice all met humane endpoints after 6 days. While mortality is not a common feature of human illness, the susceptibility allows endpoints to be easily defined for testing the effectiveness of interventions. The 6 day timepoint to death in A129 mice is also similar to that seen when the same strain of mice was been used in models of other arboviruses, including Crimean-Congo haemorrhagic fever virus (CCHFv) [49], Hazara virus [50], o’nyong-nyong virus [51, 52] and Japanese encephalitis virus [53]. Despite the severe disease and lack of a type-I interferon response, A129 mice have demonstrated protective effects with vaccines for CCHFv [54], Ross river virus [55] and chikungunya virus [56]. Therefore, by the retention of the type-II interferon response and otherwise normal immune response [57], A129 mice provide an appropriate model for investigating the adaptive immune response and performing active protective studies under stringent, frequently lethal, conditions.

When local viral loads were analysed by real-time RT-PCR, ZIKV RNA was detected in the blood, brain, ovary, spleen and liver of all A129 mice. Interestingly, viral loads in the blood, ovary, spleen and liver decreased by approximately 1-2 logs between days three and six post infection despite adverse changes in clinical symptoms, temperature and weight; however, the mean viral load in brain samples increased by over 3 logs during this period. Histological analysis showed that microscopic lesions attributable to infection with ZIKV were observed in A129 mice and these focused primarily in the brain. Lesions observed share similarities with previous studies [36] which reported necrosis with nuclear debris in Ammon’s horn (part of the hippocampus), with perivascular cuffing and astrocyte hypertrophy, in one day old Webster Swiss white mice inoculated intra-cerebrally and examined at day 7 post-challenge. In the present study, lesions generally appeared to be distributed more diffusely throughout the brain although were also seen in the hippocampus. In addition, scattered PMNs were observed. In human cases of microcephaly associated with ZIKV during pregnancy, pathological analysis has shown calcifications and other significant brain injuries [58, 59]. While similar pathological findings were not observed during this study, this is likely due to studying the acute phase of the disease in mice whereas the investigations into foetal brain abnormalities were undertaken months after the initial infection.

Whilst the main focus of attention for ZIKV is its possible links with microcephaly, as with other intrauterine infections, it is possible that the reported cases of microcephaly represent only the more severely affected children. It may be that newborns with less severe disease, affecting not only the brain but also other organs, have not yet been identified [58]; a range of aberrations in foetal development including or apart from microcephaly might occur as a result of ZIKV infection, so other possible sequelae may be possible [35]. In three children born with microcephaly that meet criteria for vertical transmission of ZIKV, funduscopic alterations in the macular region of the eyes were observed [60]. Whilst we did not study all organs in this investigation, future work may include detailed examination at ocular and other local sites.

The most significant lesions referable to infection with ZIKV in the mice were those observed in the brain. Nuclear fragmentation, perivascular cuffing by inflammatory cells, and degenerative cells in the hippocampus have been reported previously [36, 37]. It is of interest that these earlier studies involved intra-cerebral inoculation of neonatal mice; in the present study, similar lesions were observed in genetically altered, immunocompromised, adult mice challenged by subcutaneous inoculation. The microscopic lesions in the white pulp of the of A129 mice, namely large, poorly defined germinal centres, numerous apoptotic bodies and a prominent depletion in the number of mature lymphocytes could be a direct effect of virus infection or a reactive process. The marked extra-medullary haematopoiesis in the red pulp and in the liver are likely to be reactive changes to the infection process. However, immunohistochemical staining may help to clarify these possibilities by identifying whether viral antigen is present in these tissues.

Whilst no histological changes were observed in 129Sv/Ev mice challenged with ZIKV, viral RNA was detected in the blood, ovary and spleen at day 3 post-challenge and just in the ovary and spleen on day 7 post-challenge indicating that the virus entered the circulation and seeded into some of the organs. These findings suggest that despite showing no overt clinical disease, the wild-type 129Sv/Ev mice subclinically harbour ZIKV. The short timeframe of detectable viral RNA is in line with observations in humans with transient viraemia even during the acute phase of clinical disease [45, 61]. Due to the lack of overt clinical symptoms, it could be speculated that 129Sv/Ev mice might offer an option for studying the teratogenic nature of ZIKV as these mice appeared clinically healthy and challenge with ZIKV did not result in a fatal outcome.

In summary, the results we report demonstrate that A129 mice are a susceptible mouse strain to ZIKV infection which we propose as a suitable and informative *in vivo* model for the testing of vaccines and therapeutics. The challenge route and dose are similar to those found in natural infection from mosquito bite. This is the first report of a suitable murine model and can accelerate the testing of novel interventions against this pathogen declared as public health emergency of international concern.

## Acknowledgments

We would like to thank the staff of the PHE Biological Investigations Group for their assistance with the *in vivo* studies and Laura Hunter and Frances Ridley for the processing of histological samples. The views expressed in this manuscript are those of the authors and do not necessarily reflect those of the employing organisation or the funders.

